# First demonstration of the FLASH effect with ultrahigh dose-rate high-energy X-rays

**DOI:** 10.1101/2020.11.27.401869

**Authors:** Feng Gao, Yiwei Yang, Hongyu Zhu, JianXin Wang, Dexin Xiao, Zheng Zhou, Tangzhi Dai, Yu Zhang, Gang Feng, Jie Li, Binwei Lin, Gang Xie, Qi Ke, Kui Zhou, Peng Li, Xuming Shen, Hanbin Wang, Longgang Yan, Chenglong Lao, Lijun Shan, Ming Li, Yanhua Lu, Menxue Chen, Song Feng, Jianheng Zhao, Dai Wu, Xiaobo Du

**Author notes:** these authors contributed equally to this work (Feng Gao, Yiwei Yang, Hongyu Zhu).

## Abstract

Ultrahigh dose-rate (FLASH) radiotherapy has attracted immense attention because of its tumor control efficiency and healthy tissue protection during preclinical experiments with electrons, kilo-voltage X-rays, and protons. Using high-energy X-rays (HEXs) in FLASH is advantageous owing to its deep penetration, small divergence, and cost-effectiveness. This is the first report on the implementation of HEXs with FLASH (HEX-FLASH) and its corresponding application *in vivo*. With a high-current and high-energy superconducting linear accelerator, FLASH with a good dose rate and high penetration was achieved. Breast cancers artificially induced in BAL b/c mice were efficiently controlled, and normal tissues surrounding the thorax/abdomen in C57BL/6 mice were protected from radiation with HEX-FLASH. Theoretical analyses of cellular responses following HEX-FLASH irradiation were performed to interpret the experimental results and design further experiments. Thus, this study highlights the generation of HEX-FLASH for the first time and its potential in future clinical applications.

## Introduction

Cancer is the first or second leading cause of death in human beings aged less than 70 years in 91 of 172 countries and the third or fourth leading cause of death in another 22 countries^1^. Radiotherapy is an effective and one of the most widely used anti-tumor therapy methods^2^; 60%-70% of cancer patients need radiotherapy during their cancer treatment process^3^.

Though radiotherapy has a long history of more than 70 years^4^, the whole world is still trying to identify an ideal radiotherapy method, that is curable, no harm, and affordable. From 2-dimensional (2-fields penetrating to target) to intensity-modulated radiotherapy, efforts have been made to limit the area of the target by focusing majorly on the tumor and reducing radiation to adjacent organs.With the transformation of intensity-modulated radiotherapy to intensity-modulated arc therapy^5,6^, the duration and cost of radiotherapy were further minimized. With the improvement in image-guided technology, radiologists could deliver a stronger single dose to the tumor and achieve a better tumor control rate. In addition, the “Bragg peaks” of protons and heavy ions in the organism^7–9^, facilitate radiologists in delivering larger and more concentrated doses to tumors while reducing the radiation dose to surrounding normal tissues. The conventional dose-rate radiotherapy (CONV, ⩽0.1 Gy/s)^10^is a double-edged sword because although it can shrink the tumor, it can also lead to radiation-induced toxicities in normal tissues. Normal tissue complications limit the dose delivered to the tumor. Thus, the tumor cannot always be eliminated completely, and the curative ratio of radiotherapy is reduced. Therefore, eliminating tumors while protecting normal tissues has always been the main challenge of radiotherapy.

More than five decades ago, ultrahigh dose rates were noted to have some protective effects on normal tissues^11^; however, a study on isolated HeLa cells^12^revealed that the protective effects limited the clinical imagination of enhancement in cancer treatment. The effects, however, were re-recognized in 2014 where tumor cells in living mice could be suppressed by ultrahigh dose-rate radiation^13^. Since then, the new ultrahigh dose-rate (FLASH, ⩾40 Gy/s) modality has shown remarkable healthy-tissue-sparing effects in many preclinical studies without significant effects impacts on the overall treatment efficacy (called FLASH effect);its performance was superior to conventional dose-rate radiotherapy^14^. Besides its efficacy, FLASH radiotherapy was time-efficient; the total course of the treatment took only few milliseconds. Therefore, FLASH seemed to be a potential promising technique. Although the mechanisms of action remain unclear and some negative results have been reported^15^, several FLASH studies on animals^13,14,16–22^and the first human treatment in Switzerland showed satisfactory results^23^. Based on the aforementioned studies, the U.S. Food and Drug Administration approved the research device exemption for the first FLASH clinical trial^24^.

Electrons^10,13,15,19–21,25–28^, kilo-voltage (keV) X-rays^18,29^ and protons^30–32^ have been utilized in FLASH preclinical research; however, these modalities are not ideal for clinical application. Electrons and keV X-rays are usually used to treat superficial tumor sites but are not suitable for tumors situated deep inside the body, such as lung, breast, colorectal, and prostate cancers, owing to limited penetration power. FLASH protons can be used to treat deep tumors, but its high construction and operating costs discourage its use. High energy X-ray(HEX), which is most widely uesed in radiotherapy, penetrates deeply, has a small divergence, and is affordable.Therefore, it is highly suitable for clinical FLASH radiotherapy. Unfortunately, it is difficult to generate ultrahigh dose-rate HEX (HEX-FLASH); hence, it has not been used in *in vitro* or *in vivo* FLASH research, to the best of our knowledge. Therefore, HEX-FLASH is a critical component in solving the FLASH puzzle and is also an unresolved milestone for the clinical application of FLASH radiotherapy.

This study reports the first demonstration of the FLASH effect triggered by HEX on a platform called PARTER (platform for advanced radiotherapy research). In PARTER, a high-energy and high-current electron beam (6–8 MeV, >5 mA) was produced by a GaAs photocathode and superconducting linear accelerator (linac). In this study, the highest instantaneous dose rate used on experimental mice was 4 MGy/s, corresponding to a mean dose rate of over 1000 Gy/s. To verify the FLASH performance of PARTER and investigate the physiological responses of HEX-FLASH, we conducted a couple of critical mice experiments on PARTER. EMT6 tumor homografts on BALb/c mice were irradiated by HEX in FLASH and CONV to examine the tumor control efficiency of these two modalities. The thorax and abdomen of healthy C57BL/6 mice were irradiated to evaluate the FLASH effect in the normal tissue brought by HEX. Furthermore, we performed the first theoretical analysis of the HEX-FLASH effect based on the radiolytic oxygen depletion (ROD) hypothesis, and the results of this analysis provides a reference and interpretation of the FLASH effect.

## Results

### HEX-FLASH was implemented using the superconducting linac

The high-energy electron beam produced by the superconducting linac on CTFEL (Chengdu THz Free Electron Laser facility)^33,34^ was guided to bombard the rotating tungsten target on PARTER and converted to bremsstrahlung X-rays for FLASH irradiation (Fig. 1a, 1b, 1c). This procedure was essentially the same as that in the CONV radiotherapy machine; thus, the energy spectra of HEX on PARTER were similar to those of the CONV radiotherapy machine; this was confirmed by Monte Carlo computing (MCC)^35^.

**Figure 1.**
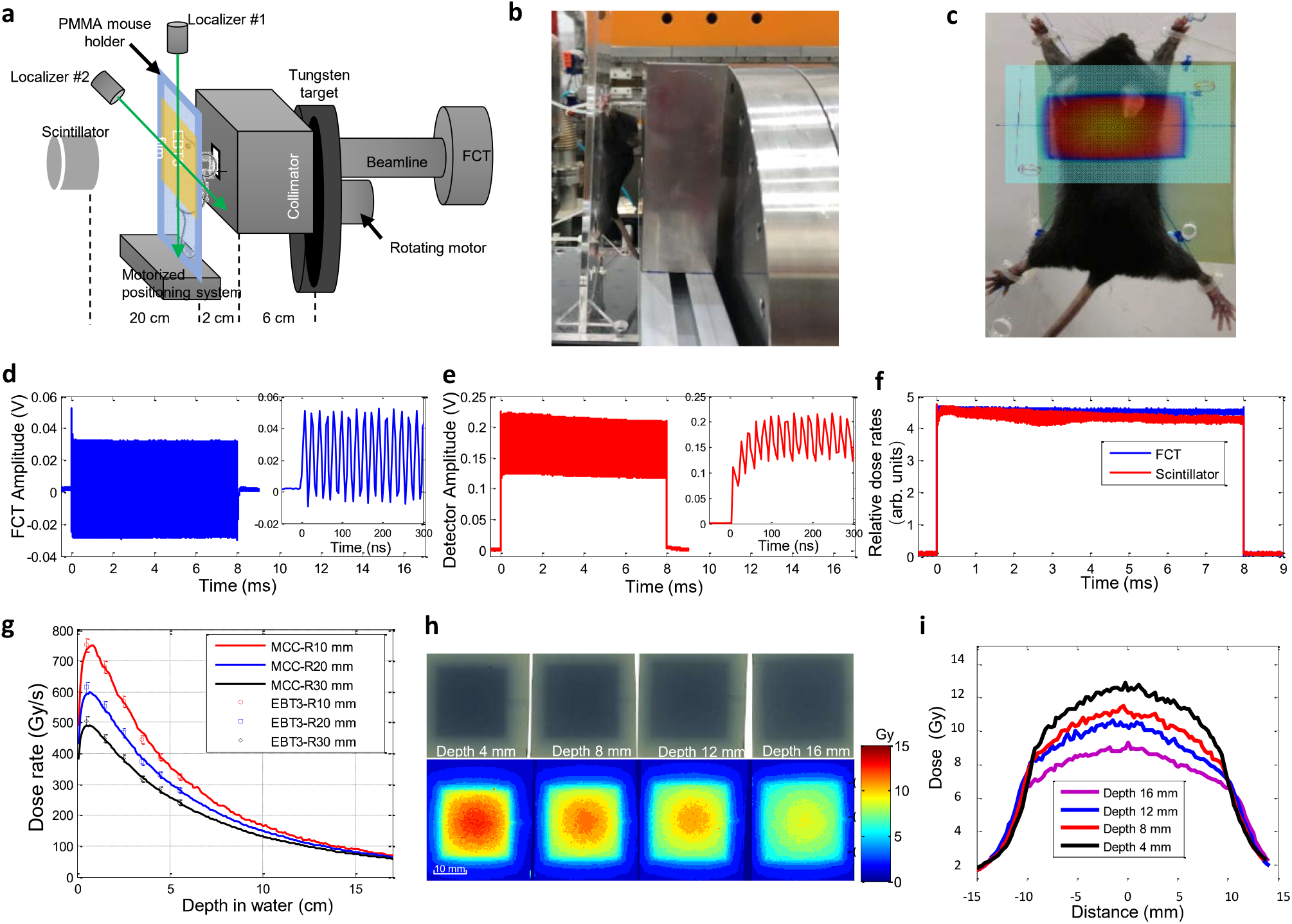
Parameters and results of the basic HEX-FLASH experiment on PARTER. **a** The schematic diagram of the HEX-FLASH experiment on PARTER. **b** A mouse was fixed on the PMMA holder while being exposed to FLASH irradiation, and **c** a EBT3 film was stuck on the backside of the holder to verify the location of the irradiation field in the mouse and to assist with dose measurement. **d** The current and pulse width of the linac measured using FCTs with a time resolution of nanoseconds, installed on the beamline, and **e, f** the time histories were compared with those obtained using the scintillator. To measure the dose rate and distribution, six EBT3 films were mounted at various depths down to up to 45 mm in the solid water phantom located at the sample position 7 cm behind the rear surface of the target chamber and irradiated using a HEX beam of 7 MeV/3.8 mA.**g** The mean dose rates within three different radii (R10, R20, and R30 mm) in the EBT3 films agree with the MCC results based on the beam current given by FCT with a discrepancy of 1% at most of the points and approximately 3% near the surface. Smaller radii show higher mean dose rates because of the center-edge dose attenuation, and the maximum mean dose rate within R10 mm, achieved at 0.5 cm depth, was approximately 750 Gy/s. The dose distribution in the phantom was measured before every biological experiment.**h** Four EBT3 films were mounted at a depth of 4–16 mm of the PMMA phantom behind the collimator window (2×2 cm) and were irradiated for 15 ms by a HEX beam of 6 MeV/5.35 mA. **i** 2D dose distribution and profiles in the x-direction in each film. HEX: high-energy X-rays; PARTER: platform for advanced radiotherapy research; PMMA: poly(methyl methacrylate); FCT: fast current transformer; MCC: Monte Carlo computing; R: radius.

The dose rate was a critical parameter that was carefully measured to monitor the implementation of FLASH. The absolute dose measured using medical radiochromic films (EBT3) showed good agreement (discrepancy <4%) with MCC results, which were based on the electron number given by the fast current transformer (FCT) device. The time structure was measured using the FCT and CeBr_3_ scintillator (Fig. 1d, 1e) and agreed with the linac character. The time histories of the dose rate given by the scintillator monitor and the FCT showed good agreement (<1%), while around 5% discrepancy was observed after approximately 1 ms when the beam power was higher than 40 kW (Fig. 1f).This discrepancy might have been induced by beam energy attenuation and has been corrected in the total dose. Both measurements and MCC results showed that in an irradiation field with a diameter of 6 cm and at a depth of over 15 cm in water, the dose rate produced by HEX on PARTER was higher than 50 Gy/s (Fig. 1g). The percentage depth dose (PDD) in the sample was adjusted by changing the beam current (1–10 mA) and energy (6–8 MV) of the superconducting linac. In our experiments, the maximum dose rate in mice was up to over 1000 Gy/s. As the depth increased, the dose rates dropped rapidly, but the dose uniformity in the irradiation field was improved.When both the collimator window and target area were set as 2 × 2 cm, the dose uniformity ratios were 1.43 and 1.25 at depths of 4 and 6 mm, respectively (Fig. 1h, 1i). MCC showed that approximately 9% of the dose deposited at the surface layer (0–2 cm) was produced by leaked high-energy electrons when the beam energy was set to 8 MeV, and it was less than 1% if the depth was over 2 cm or the beam energy was set to 6 MeV. Dose calibration was performed before every biological experiment to verify the status of PARTER and reconfirm the correlation coefficients among the dose rate, beam current, and scintillator current.

To ensure high enough dose rates, the source-surface distance (SSD) on the present PARTER platform was approximately 6–20 cm, much closer than that on the CONV radiotherapy machine (approximately 1.2 m). The closer SSD caused a higher surface dose in FLASH than in CONV, and the peak of the PDD curve in HEX-FLASH (0.7–1 cm) deviated slightly from that in CONV (approximately 1.5 cm) of the same beam energy. In the present HEX-FLASH irradiation, 2–7% of the dose delivered to mice was given by secondary electrons from the collimator window. The low beam energy corresponded to a less scattered radiation. Therefore, the recommended beam energy for HEX-FLASH in our PARTER platform is 6 MV corresponding to a maximum dose rate of approximately 800 Gy/s in mice.

### HEX-FLASH radiotherapy is as efficient as CONV radiotherapy in controlling subcutaneous homografts tumor in mice

We subcutaneously inoculated 100 ul (2 × 10^6^) EMT6 mouse breast cancer cells into the back skin of 6-week-old BAL b/c female mice. Three weeks later, we experimented with tumor-bearing mice when the tumor diameter was approximately 12–15 mm. A dose of 18 Gy in a single fraction was delivered to 15 subcutaneous homografts tumors by HEX-FLASH (900 Gy/s × 20 ms) with the linac preset to 8 MV/4.4 mA. CONV irradiation (0.1 Gy/s) was performed in the same fashion with a total dose of 15 Gy delivered to 15 tumor-bearing mice.

Observations made 63 days post-irradiation (pi) revealed that the growth rates of tumors irradiated by FLASH or CONV were slower than those observed in the control group. The slowest increase in tumor volume was achieved with FLASH radiotherapy, and the difference in tumor volume among the three groups increased with time. The tumor volume in the control group exceeded 40000 mm^3^ after 40 days pi, whereas the maximum volumes recorded in the CONV and FLASH groups were approximately 30000 and 15000 mm^3^ during the entire observation period (Fig. 2a). The difference between the CONV and FLASH groups was statistically significant (p=0.0002 in one-way ANOVA). Significant differences in volumes among the three groups were also validated with ANOVA (F = 25.14, p < 0.0001). Significant differences (p <0.005) were also observed among the survival curves of the three groups of mice (Fig. 2b); the median survival times were similar in the FLASH (55 days) and CONV (59.5 days) groups, while those in the control group were lower (35.5 days).Significantly different (p <0.005) was also observed between the survival curves of the three groups of mice(Fig. 2b), in which similar median survival time appeared in FLASH group (55 days) and CONV group (59.5 days) while control group got only 35.5 days.

**Figure 2.**
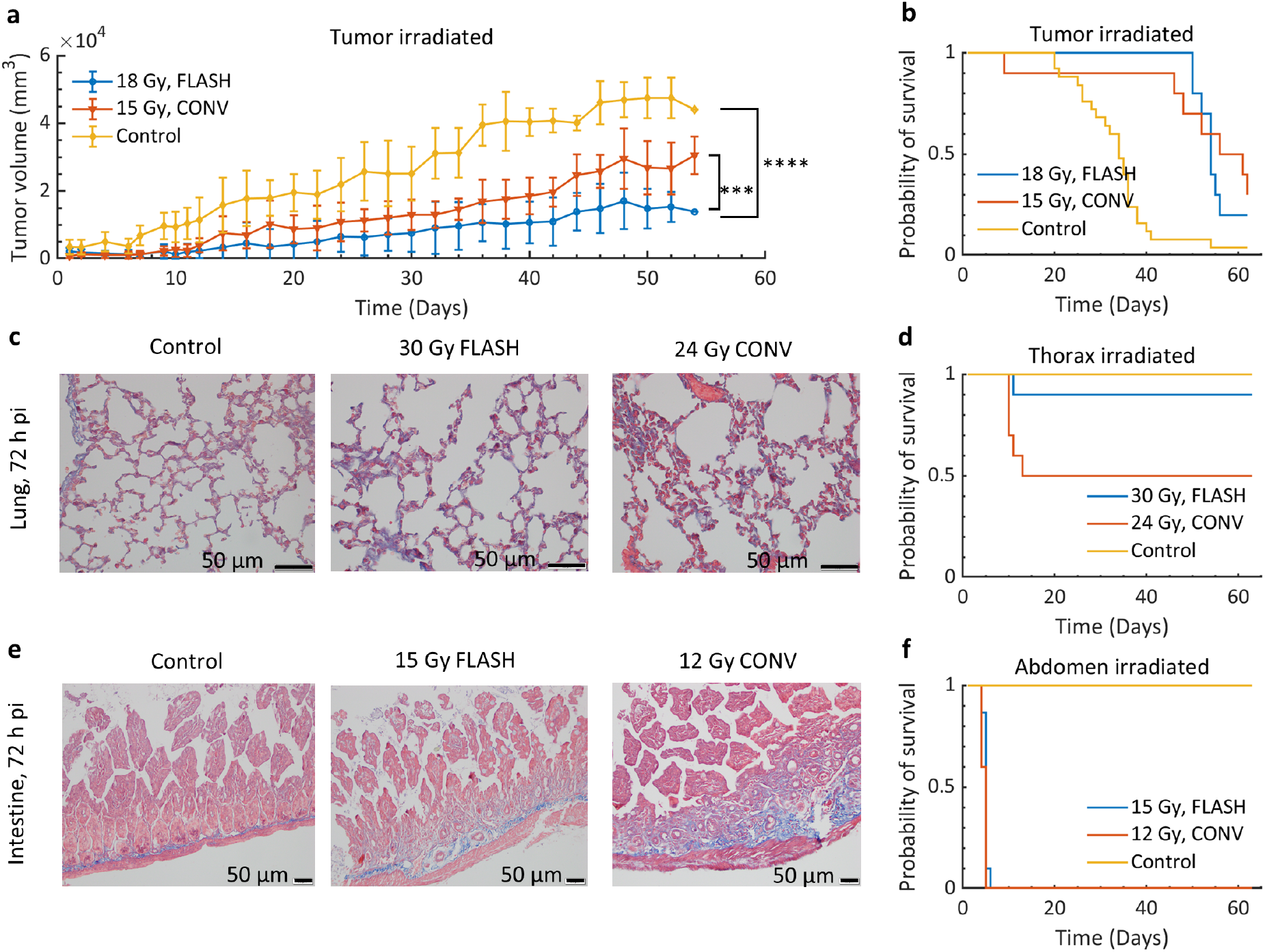
**a** Evolution of EMT6 tumor homografts in BAL b/c mice. **b** Survival curves of tumor-bearing BAL b/c mice in the control group (N=30) and after 18 Gy FLASH (N=15) and 15 Gy conventional (CONV) irradiation (N=15). **c** Masson special staining of the lung tissue of healthy C57BL/6 mice under microscope. **d** Survival curves of healthy C57BL/6 mice ofin the control group (N=19), and after 30 Gy FLASH (N=19) and 24 Gy CONV irradiation (N=20). **e** Masson special staining of the small intestine tissue of healthy C57BL/6 mice under microscope. **f** Survival curves of healthy C57BL/6 mice in the control group (N=19) and after 15 Gy FLASH (N=19) and 12 Gy CONV irradiation (N=20). pi: post-irradiation. p-values were derived with one-way repeated ANOVA: ***p < 0.001, ****p < 0.0001.

### HEX-FLASH radiotherapy leads to lesser radiation-induced damages to the lung and intestines than CONV radiotherapy

In the lung experiments, C57BL/6 mice were divided into three groups: a control group, one thorax-irradiated group receiving 30 Gy FLASH (1200 Gy/s× 25 ms), and another thorax-irradiated group receiving 24 Gy CONV (0.1 Gy/s), both in a single fraction. The dose of the CONV group was preset to a value that was 20% lower than that of the FLASH group to ensure that the actual dose received by the FLASH group was not less than that received by the CONV group even if the superconducting accelerator showed power fluctuations during FLASH irradiation (these were found to be negligible when actual measurements were made). Five mice from each group were sampled at 72 h pi for pathological examination, while the rest were retained for survival studies (they were fed). Masson special staining of the lung sections showed that alveolar structures were similar between the FLASH and control groups, whereas more disintegrated alveolar structures appeared in the lung sections of the CONV group. Collagenous fiber staining (blue-staining area) around the alveoli was more obvious in the FLASH and CONV group than in the control group. But under the microscope, we can find that the blue-staining area of the FLASH group is light and uniform, otherwise the blue-staining area of the CONV group seems to be more disorderly (Fig. 2c). The retained mice were fed for more than two months, and their survival was monitored. In thorax-irradiated mice (Fig. 2d), the survival rate was 100% in the control group, 90% in the FLASH group, and 50% in the CONV group at the end of the observation. The median survival time was not reached in the FLASH group but was 38 days in the CONV group. There was a statistically significant difference (P=0.038) in survival among the three groups. The hazard ratio (HR) was 0.19, 95% confidence intervals (CIs) were 0.035–1.010, and p was 0.0486 between the FLASH and CONV groups; therefore, the risk of death decreased by 81% in the FLASH group compared with that in the CONV group.

Similarly, in the intestine experiments,C57BL/6 mice were divided into three groups: control, FLASH, and CONV groups. The control group was common for the lung and intestine experiments. The FLASH and CONV groups received irradiation doses of 15 Gy (937 Gy/s× 16 ms) and 12 Gy (0.1 Gy/s), respectively, in a single fraction targeting the abdomen. Similar to the lung experiments, the dose of the CONV group was preset to 20% lower than that of the FLASH group, although the superconducting accelerator turned out to be stable enough. (Fig. 2e) shows that the small intestines’ basic structure still existed; however, the blue-staining area were found different in all three groups. The blue-staining area of the control group was close to muscularis propria and was very uniform and thin, while the blue-staining area of other two groups was uneven and thick, especially in the CONV group. The blue-staining area revealed the inflammatory reaction after radiotherapy led to regeneration of collagen fibers. The staining degree of FLASH group was between that of control group and the CONV group. In the CONV group, collagenous fiber staining were not only in the vicinity of the muscularis propria, but also extended to the periphery of the glandular duct (Fig. 2e). In abdomen-irradiated mice (Fig.2f), the survival rate was 100% in the control group at 63 days pi, but all the mice in the FLASH and CONV groups died at 4-5 days pi with no statistically significant difference.

To eliminate the interferences occurring owing to lethal doses and inconsistencies in the administered total doses in FLASH and CONV radiotherapies, BAL b/c mice with better radiotolerance^36^ were used in the additional experiments, in which the total doses were strictly controlled and verified. An equal dose (30 Gy) was delivered to the bilateral thoraxes of the BAL b/c mice in a single fraction of FLASH (700 Gy/s × 43 ms) and CONV (0.1 Gy/s) irradiation; the same procedure was performed on their abdomens except that the total dose was set to 12 Gy. The results showed that the survival times of mice irradiated at the bilateral thoraxes and whole abdomens were both prolonged, probably owing to the better radiotolerance of mice. The median survival times of thorax-irradiated mice in the FLASH (120 days) and CONV groups (86 days) (Fig. 3a) were statistically different (HR 0.187; 95% CI 0.044–0.803; p < 0.0001). With abdominal irradiation, the FLASH group displayed obviously better survival curves than the CONV group in which all mice lived for less than 10 days pi. (Fig. 3b). In the FLASH group, 62.5% of the mice were still alive when we stopped observation; therefore, we could not obtain a value for the median survival time. Although the survival time of mice in the FLASH group was undoubtedly higher than that in the CONV group (7 days), the difference in survival between the two groups was not statistically significant (HR 0.369; 95% CI 0.113–1.202; p=0.0735). Despite this, the survival trend of mice treated with HEX-FLASH radiotherapy was better.

**Figure 3.**
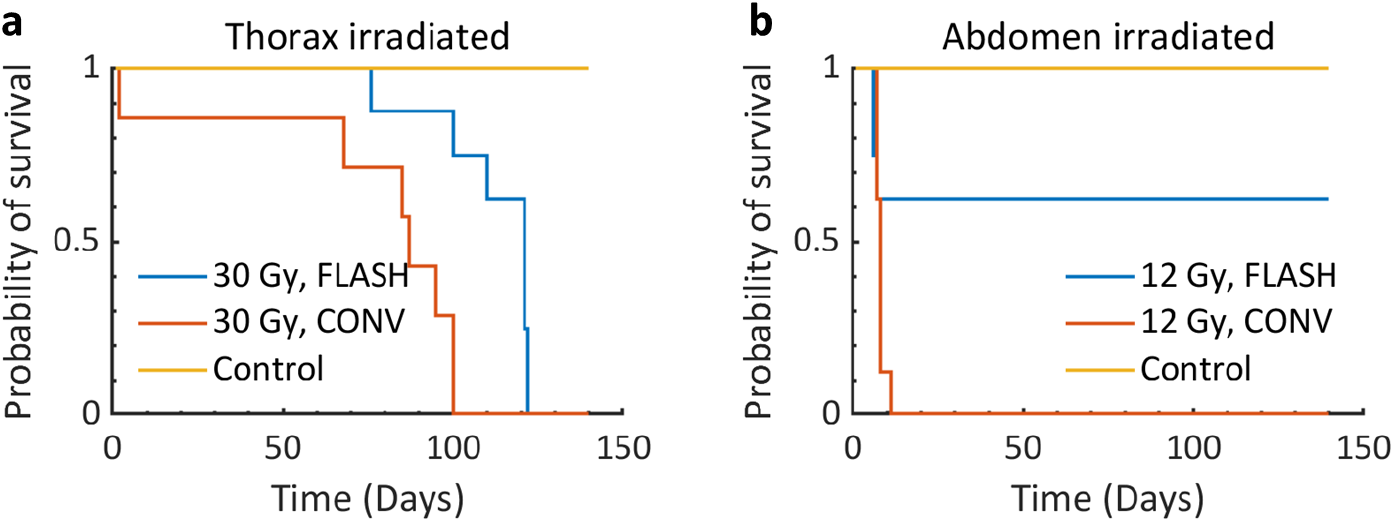
**a** Survival curves of healthy BAL b/c mice of the control group(N=8), after 30 Gy FLASH(N=8), and 30 Gy CONV(N=7) irradiation at thorax. **b** Survival curves of healthy BAL b/c mice of the control group(N=8), after 12 Gy FLASH(N=8), and 12 Gy CONV(N=8) irradiation at abdomen.

### Theoretical cellular response analysis of the FLASH effect based on the ROD hypothesis

The amount of oxygen depletion (*L*_*ROD*_) in cells (pO_2_=7.5 mmHg before irradiation) owing to HEX-FLASH irradiation was simulated to be *L*_*ROD*_ =0.196 mmHg/Gy; reduced oxygen tension in the cells led to fewer DNA damages and higher cell survival fractions (SFs). Fig. 4a presents the oxygen enhancement ratio (OER), which is defined as the ratio of the dose in anoxia to the dose in air required to achieve a given biological endpoint (e.g., a defined cell SF), as a function of oxygen tension (the scattered data points, extracted from previous experiments^37–41^, were used as references). Higher OER values at lower oxygen tensions meant that the cells were more radioresistant. When a dose of 10, 20, or 30 Gy was delivered to cells with an initial pO_2_ of 7.5 mmHg, the intracellular oxygen tension decreased, and the OER value increased from 1.41 to 1.52, 1.69, and 2.06, respectively. Fig. 4b presents the calculated SF curves of the Chinese hamster ovary (CHO) cell line irradiated with FLASH and CONV; the difference in SFs between the FLASH and CONV irradiation increased with an increase in dose. Though we considered a different cell line in the theoretical analysis, it showed consistent radioprotective effects in normal cells under FLASH irradiation.

**Figure 4.**
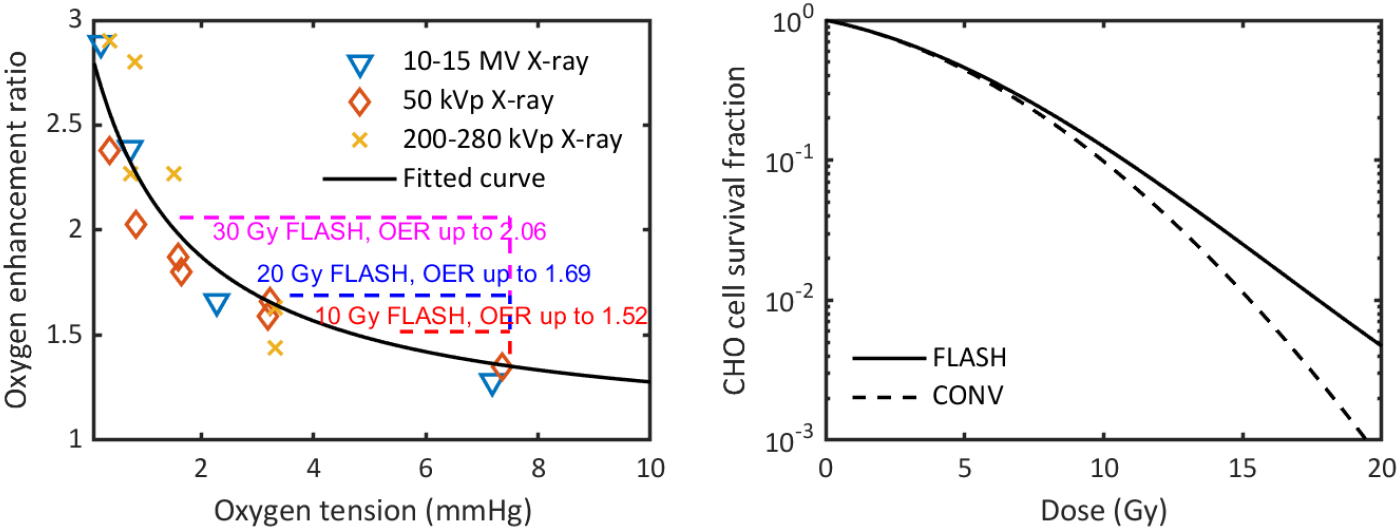
Theoretical analysis of cellular response based on the radiolytic oxygen depletion hypothesis. **a** Oxygen enhancement ratio (OER) as a function of oxygen tension (data were extracted from references^37–41^). FLASH irradiation led to rapid oxygen depletion in cells, resulting in an increase in OER values (i.e., an increase of radioresistance). **b** Survival curves of Chinese hamster ovary (CHO) cells under FLASH and CONV irradiation.

## Discussion

HEX-FLASH was implemented in PARTER without any contention according to measurement and MCC, owing to the high-current beam provided by the advanced superconducting linac at CTFEL, although the prototype is still some distance away from clinical application. As two critical parameters of HEX-FLASH, the high-energy and ultra-high dose rates have been verified by repeated measurements and careful MCC. Using HEX-FLASH, several general but critical investigations of *in vivo* physiological responses have been performed to determine whether HEX can produce FLASH effects in mice. Our results demonstrated that compared with CONV, HEX-FLASH protected the lung and intestine tissues from radiation-induced damage while efficiently controlling tumor growth.

A fundamental problem faced by our PARTER prototype was uneven dose distribution owing mainly to the absence of a flattening filter; this could be improved only by increasing the SSD. In the preclinical experiment, the SSD was set to 6–20 cm owing to the small field size for mice/cell irradiation. Moreover, according to MCC, the longest permissible SSD for the prototype was 60 cm, beyond which the dose rates would be lower than 50 Gy/s and would fall outside the FLASH range. The effective SSD could be further decreased to approximately 40 cm with the addition of a filter to equalize the dose distribution for clinical application, but this could reduce the dose rate by tens of percent. Fortunately, the performance of our PARTER prototype could satisfy the needs of the present HEX-FLASH study on animals. However, the superconducting linac at CTFEL is scheduled to be upgraded to phase II with power up to over 100 kW to generate a beam with current over three times as much as it is now; by then, the effective SSD for FLASH will be up to 100 cm, which is very close to the clinical level.

Another critically important problem was the precision of the dosimetric system; the system consisted of an EBT3 film, FCT beam monitor, and a scintillator dosimeter. The dose uncertainty given by this system was ±5%, higher than that in CONV (generally ±2%); the range of deviation was acceptable for a prototype^42^. The real-time dose rate in the sample during irradiation, computed according to the beam current given by FCT, differed from that estimated by the scintillator, especially in the early macro-pulse with high beam current. The discrepancy was because of energy attenuation in the linac. In addition, a new EBT-XD film will be used to extend its dose range to over 40 Gy. Compared with CONV radiotherapy, the ultra-short irradiation time is a significant advantage of FLASH, but it is also a big challenge for the real-time adjustment of linac characters. In the present HEX-FLASH study, the energy/current of the linac and width of the macro-pulse are preset based on calibration and computing data, but the inevitable power fluctuation of the linac during real-time irradiation leads to an actual dose different from the theoretical value (usually ±5%). Although we had considered this fluctuation in the dose design and a large sample number can counteract this effect partially in animal experiments, the interaction between the dosimetric system and linac is an important consideration for more precise experimentation and future clinical application.

In experiments determining the anti-tumor effects of HEX-FLASH, we observed significant differences in tumor volumes among the three groups (control, CONV, and FLASH groups), consistent with the results reported by Favaudon *et al*.^13^. HEX-FLASH radiotherapy showed a surprisingly excellent effect on tumor control, the tumor volume of three groups was statistically different, although the subcutaneous tumor volume was a little bit oversize when irradiated high at the time of irradiation owing to the long waiting time for the setting up of PARTER and getting permission to doing experiment (2019-nCoV outbroke in China). However, this superior effect shown by HEX-FLASH radiotherapy may probably be because of the 25% higher dose used in HEX-FLASH radiotherapy than in CONV radiotherapy. Therefore, we believe that HEX-FLASH and CONV radiotherapies have similar tumor controlling properties. This necessitates more rigorous comparison experiments in the future.

In addition, guideline of assessment for human end points in animal experiments requires tumor-bearing mice should be killed when the tumor volume exceeded 3,000 mm^3^ to alleviate the pain of experimental animals. But during the preparation period of the experiment, the 2019-nCoV outbroke in China. Considering the novelty and difficulty of this study, we put forward to the Ethics Committee that we would like to observe the survival of tumor-bearing mice at the same time. After obtaining the approval of the Ethics Committee of our hospital, we continued to feed the mice to collect more data on survival and tumor volume. After the experimental mice died, we carried out a special burial ceremony to respect and thank to them.

The tumor volume was plotted only until day 54 pi because many mice died on that day; the data after that were considered statistically insignificant. However, the survival data were not affected.Though the total dose was 25% higher in the FLASH group than in the CONV group, the FLASH group displayed protective effects on the lung tissue. This result is a powerful evidence that HEX-FLASH radiotherapy has protective effects on normal tissues. However, in intestine experiments, HEX-FLASH radiotherapy showed no protective effects. A possible explanation for this result is that different tissues have different tolerance rates to radiation; for example, the median lethal dose (LD50) of the small intestine is significantly lower than that of the lung in CONV radiotherapy^36,43,44^. Owing to poor radiotolerance of the small intestine, the dose delivered to the abdomen in the CONV group was already its lethal dose. When a 25%higher dose was delivered in the FLASH group, the mice died immediately. Thus, the protective effects of FLASH were masked.

In this study, a theoretical analysis of cellular responses following HEX-FLASH irradiation was performed based on the ROD hypothesis, which provides useful assumptions to interpret the FLASH effects. The *L*_*ROD*_ determined for HEX-FLASH was 0.196 mmHg/Gy, this allowed us to determine the cellular pO_2_ pi, and evaluate changes in oxygen-dependent radiosensitivities based on the general relationship between OER and oxygen tension (see Fig. 4a). Furthermore, the current results were calculated based on a single pulse fractionation-based dose delivery regimen, which was adopted in this study and in most of previous publications^10,14^. However, the FLASH effects triggered by different pulse fractionation regimens are also of research interest^45^. The calculation method adopted in this study can also be applied to analyze cellular responses with different fractionations; therefore, this serves as a reference for future experiment design (please see Supplementary material).

In summary, this study was conducted in three parts. First, the generation of HEX-FLASH by the PARTER system and its physical properties were confirmed. Second, the positive FLASH effects triggered by HEX were observed. Third, theoretical analyses were performed to interpret the experimental results and formulate a future experimental design. The current study provides a basis for future scientific research and clinical application of HEX in FLASH radiotherapy.

## Methods

### Irradiation devices

HEX-FLASH irradiation was performed using the PARTER platform at the CTFEL facility, Chengdu, China, in which the superconducting linac can produce 6–8 MeV electrons with an adjustable mean current of up to 10 mA and an energy spread of less than 0.2% (root mean square measured at a beam energy of 8.2 MeV). Continuous electron micro-pulses with full width at half maximum of approximately 5 ps, during which the instantaneous current is around 20 A, are produced by the linac with a period of 18.5 ns; they are then guided to bombard tungsten (3 mm in thickness) to generate HEX via bremsstrahlung. Users can freely adjust the start and end times of these continuous micro-pulses by controlling the cathode driving laser to construct a macro-pulse that is generally tens or hundreds of milliseconds.

HEX, after being collimated by purpose-built lead collimators, horizontally irradiated particular body parts of the mice when vertically fixed on a multi-hole poly(methyl methacrylate) (PMMA) plate. The SSD was adjustable from 6 cm to over 1 m, but the recommended range is 6–20 cm to meet the dose rate requirement in FLASH studies. The parameters of the linac in each irradiation were preset according to the computed dose values and calibration data before mice irradiation. For comparison, irradiation with CONV doses was performed using clinical 6 MV linac ELEKTA Precise (Elekta AB, Stockholm, Sweden) in the Mianyang Central Hospital after FLASH irradiation.

### Dosimetry

A Gafchromic^*TM*^ EBT3 radiochromic film (Ashland Inc., Covington, Kentucky, USA) was used for absolute dose measurement. Before the FLASH experiment, the EBT3 film was calibrated using a clinical 6-MV linac ELEKTA Precise (Elekta AB, Stockholm, Sweden) and a farmer ionization chamber FC65-G (IBA Dosimetry GmbH, No. 1463) associated with an IBA DOSE1^*TM*^ electrometer. The chamber was verified by the China National Institute of Measurement and Testing Technology and confirmed to be on par with international standards. The EBT3 film and ionization chamber were put at the same position in a solid water phantom and irradiated by ELEKTA Precise. The RGB value of the EBT3 film was scanned using the Epson Expression 10000XL (Seiko Epson Corporation, Japan) 24 hours after irradiation and corresponded to the dose value given by DOSE1^*TM*^. Scintillator dosimetry is still a new technology in medicine. A CeBr_3_ scintillator detector was used as a relative dosimetry system because CeBr_3_ is not an organic material. This scintillator has high X-ray detection efficiency and short raise (approximately 1 ns) and decay (approximately 20 ns) times, which make it a good technique to examine the very detailed time structure of the radiation pulse. A phototube was installed behind the scintillator to convert its optical signal into an electrical one; the electrical signal was recorded by a Tektronix MSO44 high-speed oscilloscope (Tektronix, USA) at a sampling frequency of around 1 GHz.

Two FCT devices, named FCT1 and FCT2, were installed in the accelerator for nondestructive measurement of the pulsed beam current. As a default device of the accelerator, FCT1 was calibrated in terms of the electron beam current by the CTFEL team. A Tektronix DPO4104B high-speed oscilloscope (Tektronix, USA) was used to acquire and record the FCT1 signals at a sampling frequency of around 1 GHz. FCT2 was a device dedicated for PARTER and was installed the downstream of the beamline less than one meter from the target to monitor the beam current on the tungsten target. The same Tektronix MSO44 oscilloscope used for the CeBr_3_ scintillator shared a channel to record the FCT2 signal as well; this was used only for relative electron current monitoring because it was not calibrated at the time of the experiment. All recorded signals were processed off-line by the Matlab^*R*^ 2009 program (MathWorks, USA); over 99% agreement was achieved between FCT1 and FCT2.

As a general procedure, MCC was performed using the Geant4^46^ platform to compute the dose distribution to be considered for our PARTER platform, mainly consisting of the bremsstrahlung target, collimator, and sample. As an important input of the Monte Carlo (MC) code, the accurate parameters of the electron beam were given by the advanced superconducting linac and FCT devices. These accurate structures and parameters could validate the exactness and reliability of the MCC, which were already verified through several calibration experiments using EBT3 film for comparison.

### Theoretical cellular response analysis after HEX-FLASH irradiation

Theoretical analyses of the FLASH effects were performed with MCC and previously developed radiobiological models to interpret the experimental results. Theoretical analyses were based on the ROD hypothesis^10,47,48^, a leading hypothesis on the radioprotective effects experienced by normal tissues exposed to FLASH irradiation. The ROD hypothesis considers that oxygen molecules get rapidly depleted during FLASH irradiation and create a local radiobiological hypoxic environment, thus making cells less radiosensitive. Pratx and Kapp proposed that hypoxic stem cells may be related to the radioprotection from ROD associated with the FLASH effect^48,49^. Therefore, hypoxic cells (with an oxygen tension pO_2_ = 7.5 mmHg) were considered for the analyses of cellular response.

The nanodosimetry Monte Carlo simulation code (NASIC) was adopted for simulating the cellular responses after CTFEL HEX-FLASH irradiation. NASIC is a MC track structure tool that supports simulations of physical and chemical reactions induced by ionization irradiations and analyses of subsequent radiobiological responses, including DNA damage and cell SFs under CONV^50–53^ and FLASH^54^. The detailed description of the MC simulation method and analysis model for different biological endpoints can be found in the cited references^54,55^.

DNA double-strand breaks (DSBs) induced by ionization radiation were assumed as the initial and most lethal lesions in the cell nucleus and were positively related to cell death after irradiation. In the NASIC MC simulation, it was assumed that direct DNA damages were induced by physical reactions between incident particles and the DNA structure, and indirect DNA damages resulted from reactions between molecular oxygen (O_2_) and DNA-derived radicals, which were produced by reactions between DNA structures and hydroxyl radicals (·OH). The yield of DNA double-strand breaks changed with oxygen tension (pO_2_) in cells and resulted in different cell SFs.

In this study, the initial DNA damage information and amount of oxygen depletion (*L*_*ROD*_) under different fractionation schemes were simulated with NASIC. The dynamic evolution of pO_2_ in cells and changes of final DNA damage yield as well as cell SFs were assessed by following the method described in the references^54,55^; the cell line used in the analyses was CHO cells.

### Animal experiment and ethics statement

We subcutaneously inoculated EMT6 mouse breast cancer cells into the back skin of 6-week-old BAL b/c female mice. We experimented with tumor-bearing mice when the tumor diameter was approximately 12~15 mm. The tumor-bearing mice were divided into three groups: a control group, a group irradiated by FLASH (1000 Gy/s) with 18 Gy/1F, and a group irradiated by CONV (0.1 Gy/s) with 15 Gy/1F. The radiation field was square shaped with sides of length 1.5 cm. After the mice were irradiated, we observed the mice every day and recorded the long and short diameters of the tumors every other day. We observed tumor-bearing mice for 63 days. The volume was calculated using the following formula: V = 0.52 long diameter short diameter^2^. After calculating the volume of the tumor in each mouse, we averaged the tumor volumes and drew a line chart. We recorded the number of mice alive every day and finally drew a survival curve of the tumor-bearing mice.

Six-week-old C57BL/6 female mice were chosen for our normal tissue radiation experiments. For lung experiments, the C57BL/6 female mice were divided into three groups: control, FLASH-irradiated, and CONV-irradiated groups. We irradiated the bilateral thoraxes of the mice in the FLASH and CONV groups and confirmed that the upper limit of the thorax was the auricle’s lower edge in natural condition; the lower limit was 2 cm below the upper limit. Nineteen mice and twenty mice from each group were exposed to either FLASH or CONV radiotherapy. In the FLASH group, the mice were irradiated with HEXs having a dose rate of 1200 Gy/s; the total dose was 30 Gy. In the CONV group, the mice were irradiated with X-rays having a dose rate of 0.1 Gy/s, with the total dose being 24 Gy.

For intestine experiments, the C57BL/6 female mice were divided into three groups: control (the same group as mentioned above), FLASH-irradiated, and CONV-irradiated groups. The irradiation field was 2 × 4 cm^2^ square. The upper edge was defined as the lower part of the thorax (2 cm below the lower edge of the auricle) and the lower edge was 4 cm below the bottom of the thorax. Nineteen and twenty mice from each group were exposed to either FLASH or CONV radiotherapy. In the FLASH group, the mice were irradiated with HEXs having a dose rate of 937 Gy/s; the total dose was 15 Gy. In the CONV group, the mice were irradiated with X-rays having a dose rate of 0.1 Gy/s; the total dose was 12 Gy.

After the mice were irradiated, we observed their survival for 63 days. Lung and small intestine tissues were obtained for pathological examination 72 hours pi. We used a microscope to observe the lung tissue sections stained with Masson special staining and magnified by 200 or 400 times. At the end of the observation, we compiled the survival data into a survival curve. In the second experiment, to eliminate the interferences occurring owing to inconsistencies in the administered total doses in FLASH and CONV radiotherapies, we followed the exact protocols mentioned above. The differences between the two experiments on normal tissue were the different strain of mice and dose delivered to targets. We chose 6-week-old BAL b/c female mice and strictly verified that an equal dose (30 Gy) was delivered to the bilateral thoraxes of the BAL b/c mice in a single fraction of FLASH (700 Gy/s × 43 ms) and CONV (0.1 Gy/s) irradiation, the same procedure was performed on their abdomens except that the total dose was set to 12 Gy. All the mice mentioned in this paper were purchased from Sichuan University, China. Before the experiments were conducted, approval regarding animal ethics was obtained from the Animal Ethics Committee of Mianyang Central Hospital (approval number: P2020032). For animal experiment we followed guideline of assessment for human end points in animal experiment published in 2018 in China. Owing to the huge contribution from experimental tumor-bearing mice in our experiment, we completed our observation of tumor volume and survival. We carried out a special memorial service to thank them.

### Statistics analyses

In our research, we calculated the average tumor volume in the tumor-bearing mice and overall survival of mice irradiated at different dose rates. Statistical analysis was performed using GraphPad Prism v.8.4.0 (GraphPad, La Jolla, CA) software. The balance between the FLASH and CONV groups was examined using the chi-square test or independent-sample t-test. Single factor analysis of variance (ANOVA) was used to analyze the changes in the mean tumor volumes of tumor-bearing mice in the three groups. Results were expressed as mean values ± SD. Survival curves were compared using the Kaplan-Meier method with a log-rank test. All statistical tests were conducted at a significance level of 5%, and a two-tailed p-value <0.05 indicated a statistically significant difference; 95% CIs were also calculated.

### Data availability

The authors declare that the data supporting the findings of this study are available within the paper and in the Supplementary Information file. All other data are available from the authors upon reasonable request.

## Acknowledgements

This work was financially supported by the National Natural Science Foundation of China (NSFC grant nos. 11975218, 11905210, 11805192, 12005211, and 11605190) and Innovation Foundation of CAEP (grant nos. CX2019036 and CX2019037).

## Author contributions statement

Y.Y., X.D., F.G., and D.W. conceived the idea; X.D., F.G., Y.Y., D.W. and H.Z. designed the study; J.W., D.X., Z.Z., T.D.,Y.Z.,G.F.,J.L.,B.L., K.Z., P.L., X.S., H.W., L.Y., C.L., L.S., M.L., S.F., Y.L., M.C. and J.Z. conducted the experiments; F.G. and Y.Y. performed the data analysis; H.Z. carried out the simulations; F.G., Y.Y., H.Z., D.W. and X.D. interpreted the results; G.X. and Q.K. observed the morphological changes in mice tissues under the microscope; Y.Y., F.G., H.Z., D.W., Z.Z. and X.D. wrote the manuscript; X.D. and D.W. supervised the investigation. All authors reviewed the manuscript.

## Competing interests

The authors declare no competing interests.

## Additional information

To include, in this order: **Accession codes** (where applicable); **Competing interests** (mandatory statement).

The corresponding author is responsible for submitting a competing interests statement on behalf of all authors of the paper. This statement must be included in the submitted article file.

## Supplementary

### Theoretic analysis of the FLASH effect of different pulse fractionations

Three pulse fractionation regimens, i.e., a total dose of 20 Gy delivered to the target tissue in 1, 2, or 4 fractions, were considered for analysis. Fig. S1a shows the rapid oxygen depletion due to FLASH irradiation and the recovery of pO_2_ due to oxygen diffusion from blood vessels after each fraction. Fig. S1b shows that the pO_2_ values drop to the lowest level within the time scale of 40 *μ*s, the low pO_2_ level can hold for approximately 10 ms, and the oxygen tension in tissues is generally fully recovered within 1 second.

**Figure S1.**
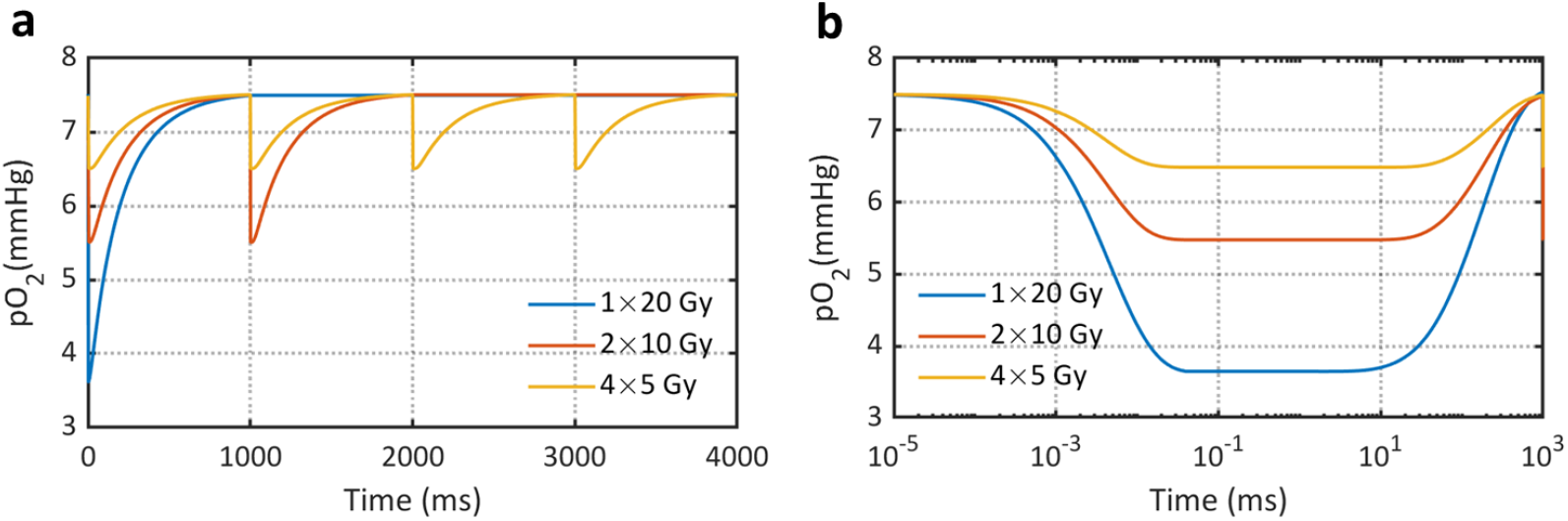
Dynamic change of pO_2_ in cells that were exposed to HEX FLASH radiation with a total dose of 20 Gy delivered in 1, 2, and 4 fractions. **a** Dynamic change of pO_2_ in cells after each fraction; **b** Semi-log plot of dynamic change of pO_2_ in cells after the first fraction.

Different fractionations of FLASH irradiation resulted in differences of pO_2_ evolution after each pulse, the pO_2_ value after each pulse dominates the indirect DNA damage yield as well as cell SFs. Table S1 shows the cellular responses of CHO cells toward different FLASH irradiation schedules. The pO_2_ value after the irradiation of 20 Gy in 1 fraction of was 45.1% lower than that after the irradiation of 20 Gy delivered in 4 fractions, and this further led to 13.5% less indirect DNA damages and 75.6% higher cell SF.

**Table S1.** The pO_2_ value after each pulse, indirect DNA damage yield, and cell SFs of CHO cells (pO_2_ =7.5 mmHg) were exposed to CTFEL X-ray FLASH radiation with a total dose of 20 Gy delivered in 1, 2, and 4 fractions.

| Fractionation | The pO<sub>2</sub> value after each pulse (mmHg) | Indirect DNA damage yield (Gbp<sup>-1</sup>Gyp<sup>-1</sup>) | Cell SFs |
| --- | --- | --- | --- |
| 1 × 20 Gy | 3.58 | 4.82 | 0.47% |
| 2 × 10 Gy | 5.54 | 5.37 | 0.16% |
| 4 × 5 Gy | 6.52 | 5.57 | 0.10% |
| Max. rel. diff.* | 45.1% | 13.5% | 75.6% |

\* Max. rel. diff: maximum relative difference was calculated with (1-min./max.) × 100%

## Notes

### Competing Interest Statement

The authors have declared no competing interest.

### Summary of Updates

Section on HEX-FLASH radiotherapy leads to lesser radiation-induced damages to the lung and intestines than CONV radiotherapy updated; Figure 2 and Figure 2 revised

